# *Plasmodium berghei* K13 Mutations Mediate *In Vivo* Artemisinin Resistance That is Reversed by Proteasome Inhibition

**DOI:** 10.1101/2020.08.24.256842

**Authors:** Nelson V. Simwela, Barbara H. Stokes, Dana Aghabi, Matt Bogyo, David A. Fidock, Andrew P. Waters

## Abstract

The recent emergence of *Plasmodium falciparum* (PF) parasite resistance to the first line antimalarial drug artemisinin is of particular concern. Artemisinin resistance is primarily driven by mutations in the PF K13 protein, which enhance survival of early ring stage parasites treated with the artemisinin active metabolite dihydroartemisinin *in vitro* and associate with delayed parasite clearance *in vivo*. However, association of K13 mutations with *in vivo* artemisinin resistance has been problematic due to the absence of a tractable model. Herein, we have employed CRISPR/Cas9 genome editing to engineer selected orthologous PF K13 mutations into the *K13* gene of an artemisinin-sensitive, *P*. *berghei* (PB) rodent model of malaria. Introduction of the orthologous PF K13 F446I, M476I, Y493H and R539T mutations into PB K13 produced gene-edited parasites with reduced susceptibility to dihydroartemisinin in the standard 24-hour *in vitro* assay and increased survival in an adapted *in vitro* ring-stage survival assay. Mutant PB K13 parasites also displayed delayed clearance *in vivo* upon treatment with artesunate and achieved faster recrudescence upon treatment with artemisinin. Orthologous C580Y and I543T mutations could not be introduced into PB while the equivalent of the M476I and R539T mutations resulted in significant growth defects. Furthermore, a *Plasmodium*-selective proteasome inhibitor strongly synergized dihydroartemisinin action in these PB K13 mutant lines, providing further evidence that the proteasome can be targeted to overcome ART resistance. Taken together, our work provides clear experimental evidence for the involvement of K13 polymorphisms in mediating susceptibility to artemisinins *in vitro*, and most importantly under *in vivo* conditions.

**IMPORTANCE:** Recent successes in malaria control have been seriously threatened by the emergence of *Plasmodium falciparum* parasite resistance to the frontline artemisinin drugs in Southeast Asia. *P*. *falciparum* artemisinin resistance is associated with mutations in the parasite K13 protein, which associates with a delay in the time required to clear the parasites upon treatment with the drug. Gene editing technologies have been used to validate the role of several candidate K13 mutations in mediating *P*. *falciparum* artemisinin resistance *in vitro* under laboratory conditions. Nonetheless, the causal role of these mutations under *in vivo* conditions has been a matter of debate. Here, we have used CRISPR/Cas9 gene editing to introduce K13 mutations associated with artemisinin resistance into the related rodent-infecting parasite, *P*. *berghei*. Phenotyping of these *P*. *berghei* K13 mutant parasites provides evidence of their role in mediating artemisinin resistance *in vivo*, which supports *in vitro* artemisinin resistance observations. However, we were unable to introduce some of the *P*. *falciparum* K13 mutations (C580Y, I543T) into the corresponding amino acid residues, while other introduced mutations (M476I, R539T equivalents) carried a pronounced fitness cost. Our study provides evidence of a clear causal role of K13 mutations in modulating susceptibility to artemisinins *in vitro* and *in vivo* using the well-characterized *P*. *berghei* model. We also show that inhibition of the *P*. *berghei* proteasome offsets parasite resistance to artemisinins in these mutant lines.

## INTRODUCTION

Artemisinin (ART)-based combination therapies (ACTs) have been at the forefront of globally coordinated efforts to drive down the burden of malaria. A pharmacodynamic hallmark of ARTs and their derivatives is that they are highly active and fast acting against blood stages of malaria parasites. These drugs can achieve up to 10,000 fold parasite reductions in the first replication cycle upon drug exposure (1). Such is the effectiveness of ARTs that recently reported reductions in malaria morbidity and mortality are, indeed, partly attributed to ACTs (2). The use of ARTs in combination therapies originated from early clinical trials which showed that despite achieving faster parasite clearance, ART monotherapies resulted in recrudescence rates of up to 40% (3). ACTs deliver a pharmacological cure by taking advantage of ARTs to rapidly clear the parasite biomass in the early days of treatment, while relying on the partner drug to eliminate residual parasites (4). So far, ACTs remain highly effective in Sub-Saharan Africa, the region that harbors the highest disease burden, with efficacy rates of >98% (2). Nevertheless, ACTs have been threatened by the emergence of *Plasmodium falciparum* (PF) resistance to ARTs in Southeast Asia (SEA), and resistance has the potential to spread to other malaria-endemic regions, as has been a historical trend with earlier first-line antimalarial drugs (2, 5, 6). Indeed, K13 variants that are able to mediate ART resistance *in vitro* have recently been identified in PF parasites in French Guiana and in Rwanda (7, 8). Moreover, recent aggressive expansion of a parasite lineage carrying the genetic determinants of resistance to both ART derivatives and the ACT partner drug piperaquine has been reported across SEA, resulting in a dramatic loss of clinical efficacy (9, 10).

Clinically, PF resistance to ARTs manifests as reduced *in vivo* parasite clearance upon treatment with ACTs or ART monotherapies (2, 5, 6). These clearance rates are based on the WWARN parasite clearance estimator (11), which quantifies relative resistance by estimating parasitemia lag phases and clearance half-lives upon treatment with artesunate (AS) or ACTs. This involves *in vivo* quantification of viable parasitemia (in patients) upon treatment with AS (2-4 mg/kg/day) or ACTs at specified time intervals, and subsequent calculation of parasite densities as a function of time (11). The parasite clearance estimator has been used to generate substantial baseline data that classify ART resistance as parasite clearance half-lives >5.5 hours and ART sensitivity as parasite clearance half-lives <3 hours (12, 13). However, interpretation of clearance half-lives can be confounded by differences in initial parasite biomass, the efficacy of the partner drug, and the level of host immunity (12, 14). Moreover, this *in vivo* phenotype does not correlate with decreased susceptibility to dihydroartemisinin (DHA) in standard growth inhibition assays where PF parasites (which have an ∼48 hour intra-erythrocytic developmental cycle) are exposed to the drug for a total of 72 hours (5, 15, 16). The ring-stage survival assay (RSA), where highly synchronized early ring-stage parasites (0-3 hours post invasion) are exposed for a short period of time (3-6 hours) to DHA (at the pharmacologically relevant concentration of 700 nM) provides an improved correlate for the *in vivo* delayed parasite clearance phenotype, and has been the principal *in vitro* assay for determining PF resistance to ARTs (17). At the genetic level, polymorphisms in the PF K13 propeller domain have been strongly associated with ACT treatment failure (16, 18) and also correlate with delayed parasite clearance *in vivo* and increased parasite survival *in vitro* in RSAs (19-21). Reverse genetic approaches have been successfully used to show that the PF K13 mutations M476I, R539T, I543T, Y493H and C580Y can confer DHA resistance *in vitro*, as defined by >1% survival in RSAs (22, 23). However, the parasite genetic background as well as underlying polymorphisms in drug resistance determinants such as *pfcrt* and *mdr2* may play a role either by modulating different levels of susceptibility to DHA or providing a suitable biological landscape upon which these K13 mutations are more likely to arise (19, 22).

ART resistance as typified by the “delayed clearance phenotype” is, however, still classified as “partial resistance”, mostly because most patients with parasites harbouring the phenotype effectively clear the infection when an effective partner drug is used or duration of monotherapy is extended (4). ART “partial resistance” is, therefore, confirmed or suspected when patients carry parasites with certain K13 mutations, display a parasite clearance half-life > 5.5 hours, or are microscopically smear positive on day three after initiation of treatment (2, 4). The full extent to which these parameters predict subsequent ACT treatment failure or define ART resistance remains an area of continuing debate (24-29). The definition of ART resistance in these contexts would thus benefit from experimentally accessible *in vivo* models that would help interrogate ART parasite susceptibility parameters including clearance half-lives, recrudescence rates, and treatment failures. Such models would allow for a genetic dissection of the role of K13 mutations in mediating resistance *in vivo* in the absence of confounding factors such as secondary genetic factors and/or host factors (19, 22). Currently, the K13 C580Y polymorphism is the most prevalent and dominant ART-resistant mutation in SEA (6, 30). A recent genetic cross of the K13 C580Y ART-resistant line with an *Aotus* monkey-infecting PF strain provided evidence, in this non-human primate model, that parasites carrying the C580Y mutation could display increased survival in *in vitro* RSAs with no accompanying *in vivo* ART resistance (31).

Moreover, *P*. *falciparum* drug resistance mutations are known to often associate with significant fitness costs that limits the prevalence and eventual propagation of resistance-conferring alleles in natural infections. For example, mutations in chloroquine (CQ) resistance transporter (*pfcrt)* that modulate resistance to CQ massively expanded when CQ was in use in the 1970s but eventually were outcompeted and replaced with parasites carrying wild-type alleles in African high-transmission settings following withdrawal of CQ use (32, 33). Similarly, PF K13 mutations have been shown to carry *in vitro* fitness costs, however, the degree to which a given mutation is detrimental for growth seems to depend on the parasite genetic background (34). Relative to other K13 mutations, PF R539T and I543T mutant parasites that are associated with the highest RSA survival rates (20, 22) and most significant delays in parasite clearance (35) also carried the most pronounced fitness costs (34). Intriguingly, the most prevalent SEA K13 mutation, C580Y, was fitness neutral *in vitro* when gene edited into recent Cambodian clinical isolates, whereas it displayed a significant growth defect when introduced into ART-susceptible parasites isolated before ARTs were widely deployed (34, 36). Recently, it has been demonstrated that PF K13 localizes to the parasite cytostomes and other intracellular vesicles and plays a role in parasite hemoglobin endocytosis and trafficking to the lysosome-like digestive vacuole (37-39). K13 mutations are thought to lead to a partial loss of protein function, which subsequently impairs hemoglobin endocytic uptake, thereby lessening ART activation and conferring ART resistance (37). This has pointed towards a K13-mediated hemoglobin-centric mechanism of ART resistance, which could be possibly shared with other drugs such as CQ that act by binding to heme moieties in the digestive vacuole, following cytostome-mediated hemoglobin endocytosis (38, 40-42). Of note, mutant K13-mediated ART resistance phenotypes are associated with upregulated cellular stress responses, which can be targeted by selective inhibition of the parasite 26S proteasome (43, 44).

Here, we report the *in vitro* and *in vivo* phenotypes of orthologous PF K13 mutations that were gene-edited into an *in vivo* rodent model of malaria, *P*. *berghei* (PB). We profiled the fitness of these PB K13-mutant parasites relative to their isogenic wild-type counterparts as well as their sensitivity to combinations of DHA and proteasome inhibitors. Our data provide evidence that K13 mutations are causal for reduced susceptibility to ARTs in an *in vivo* model and link these mutations to *in vitro* and *ex vivo* phenotypes. Our findings also demonstrate that inhibition of the *Plasmodium* proteasome is an effective strategy to restore ART action in resistant parasites that survive treatment with ART alone.

## RESULTS

### CRISPR/Cas9-mediated introduction of PB orthologous K13 mutations and *in vivo* mutant enrichment by AS

To generate PB mutant parasites carrying orthologous PF K13 mutations, we attempted to introduce PB equivalents of five PF K13 mutations (M476I, Y493H, R539T, I543T and C580Y) that by reverse genetics have been previously shown to confer enhanced PF survival in *in vitro* RSAs (22). We also introduced the equivalent of the F446I mutation that is predominant in Southern China along the Myanmar border (6). These mutations are all validated determinants of reduced PF susceptibility to ARTs (4). Structural homology modelling revealed that PB and PF K13 (PBANKA_1356700 and PF3D7_1343700, respectively) are highly conserved (∼84% sequence identity overall) at the C-terminal propeller domain, especially where resistance-conferring mutations localize (**Fig. 1A**). PB K13 carries 12 extra amino acids, resulting in 738 amino acids for PB compared to 726 for PF. However, modelling suggests that the extra amino acids in PB do not change the overall propeller structure of K13 or the amino acid identity at the orthologous positions of the mutations examined in this study (**Fig. 1A; Fig. S1A, S1B**). Using a CRISPR/Cas9 system (**Fig. S2A**) (40), we designed Cas9 plasmids carrying single guide RNAs (sgRNAs) to target the PB K13 locus with corresponding homology repair templates. The repair templates carried the mutations of interest as well as silent mutations that inactivated the protospacer adjacent motif (PAMs) and introduced restriction sites for restriction fragment length polymorphism (RFLP) analyses (**Table S1**). Electroporation of the plasmids pG1004 (C592Y), pG1005 (I555T) and pG1006 (R551T) into the K13 wild-type PB 1804cl1 line yielded edited parasites (G2022^C592Y.1*^, G2023^C592Y.2*^, G2024^I555T*^ and G2025^R551T*^) with calculated 13.4%, 18.5%, 7.7% and 30.0% efficiencies respectively by RFLP analysis (**Fig. S2B; Table S1**). Intriguingly, bulk DNA sequencing of these transformed parasites revealed that only the G2025^R551*^ line carried sequence traces for the R551T amino acid substitution and accompanying silent mutations (**Fig. S3C**) while the rest had traces only of the silent mutations (**Fig. S3A, S3B**). These data suggested that the C592Y and I555T mutations either result in extremely slow growing parasites or are entirely lethal in PB. We attempted to clone the G2025^R551T*^ line by limiting dilution, but this could not be achieved, possibly due to the low mutant population (30.0 %) combined with a potentially slow growth rate of the mutants compared to wild-type parasites.

**Fig. 1.**
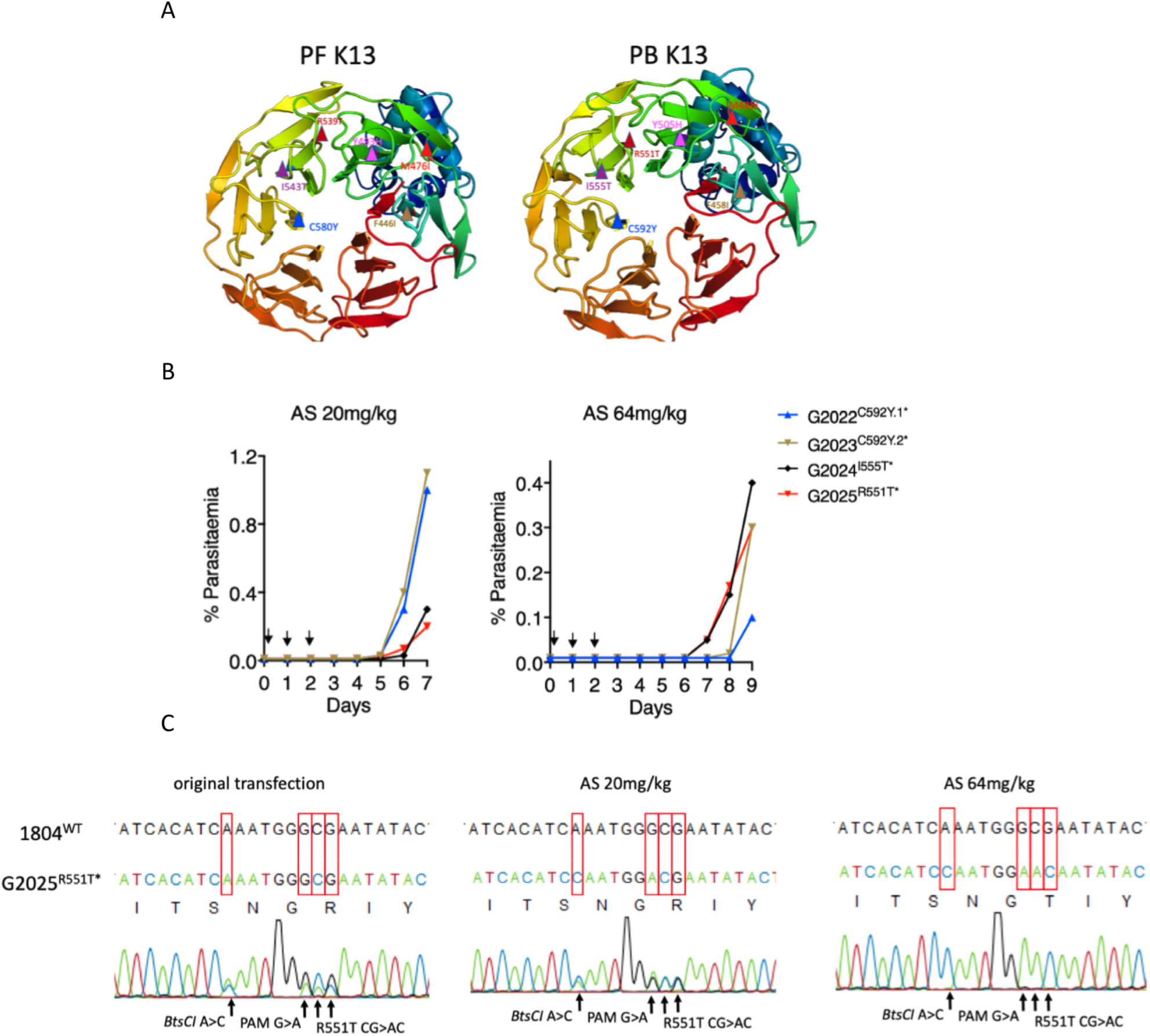
Introduction of orthologous K13 nucleotide substitutions in PB. **A**. 3D homology model of PF (PF3D7_1343700) and PB (PBANKA_1356700) K13 for amino acid residues 350-726 and 362-738, respectively. PF K13 mutation sites (F446I, M476I, Y493H, R539T, I543T and C592Y) are indicated in the structure on the left and PB orthologous mutation sites are modelled on the right. Models were created in SwissModel using PDB template 4zgc.1.A. Structures were visualized and annotated using pyMol 2.3. **B**. Parasitemia growth curves monitoring recrudescence of the G2022, G2023, G2024 and G2025 lines upon artesunate (AS) challenge. Mice were infected with 2×10^7^ parasites by IP injection on day 0. Treatment with AS was commenced ∼3 hours post infection by IP and was continued for three consecutive days as indicated by arrows. Parasitemia was monitored microscopically until recrudescence was observed. Mice were bled when the parasitemia was less than 1.5% to minimize competition from wild-type parasites in case mutants carried growth defects. **C**. Sanger sequencing of bulk DNA from the G2025 R551T line showing selective enrichment of this mutation upon AS treatment at 20 or 64 mg/kg. Enrichment of this mutation was also observed in the RFLP analysis (**Fig. S2B**).

In earlier efforts to introduce UBP-1 mutations in PB, we found that pre-emptive drug pressure to which the engineered mutation is anticipated to confer protective advantage can selectively enrich for the mutant in a mixed, transfected parasite population even when the mutant population is <1% in the mixture (40). Using this approach, we subjected a larger inoculum (2 x 10^7^) of the G2022^C592Y.1*^, G2023^C592Y.2*^, G2024^I555T*^ and G2025^R551T*^ lines to AS at 20 or 64 mg/kg to see if any enrichment in the recrudescent parasite populations could be achieved (**Fig. 1B**). Indeed, AS at both 20 and 64 mg/kg specifically enriched the R551T mutant population in the G2025^R551T*^ line from 30.0% in the initial transfection to 49.7% at AS 20 mg/kg and >99% at 64 mg/kg (**Fig. 1C, S2B; Table S1**). In contrast, apart from a minor enrichment that was observed for the G2024^I555T*^ line, no useful enrichments in both the G2022^C592Y.1*^ and G2023^C592Y.2*^ lines were observed by RFLP at either concentration of AS (**Fig. S2B; Table S1**). Furthermore, no I555T or C592Y amino acid substitution traces could be seen after population-level DNA sequencing of these lines. These data further supported the relative non-viability of PB parasites bearing K13 C592Y and I555T mutations. In agreement with the above observations, further attempts to introduce the C592Y mutation using a different sgRNA and or different codons for the tyrosine residue in the donor template (TAT or TAC) were also unsuccessful. We did, however, observe >90% editing efficiency when introducing only silent mutations that maintained the C592C wild-type genotype in the donor template (**Fig. S3E, S3F; Table S1**). This, plus other unsuccessful attempts to generate the I555T mutant further implies that these two K13 mutations are not viable in PB. Meanwhile, transfection of the PB 1804cl1 line with pG983 (F458I), pG984 (Y505H) and pG1008 (M488I) successfully introduced these mutations in PB K13, yielding the G1957^F458I*^, G1979^Y505H*^ and G1989^M488I*^ lines with >93% efficiencies as confirmed by RFLP analysis (**Fig. S2C; Table S1**) as well as population level DNA sequencing (**Fig. S3G, S3H, S3I**). These three lines (G1957^F458I*^, G1979^Y505H*^, G1989^M488I*^) and the G2025^R551T*^ AS 64 mg/kg challenged line were all cloned by limiting dilution. Mutations were further confirmed by RFLP analysis (**Fig. S3D**) and sequencing. The V2721F UBP-1 mutant line, which we previously found to mediate reduced susceptibility to ARTs in PB (40), was also generated in the 1804cl1 background and cloned (**Table S1**).

### PB K13 mutants display reduced susceptibility to DHA in 24-hour assays and increased survival in PB-adapted RSAs

Unlike in PF, PB can only be maintained in one blood stage cycle *in vitro*, which restricts drug susceptibility assays to one 24-hour developmental cycle. Drug susceptibility readouts are, therefore, based on single-generation flow cytometry quantification of schizont maturation (40, 45, 46). Using this approach, we aimed to characterize the DHA dose-response profiles of the PB K13 mutants as compared to wild-type parasites or to a previously reported UBP-1 mutant with reduced ART susceptibility (40). Interestingly, in contrast with the equivalent PF K13 mutants, PB M488I, R551T and Y505H K13-mutant parasites displayed reduced susceptibility to DHA in standard growth inhibition assays with 3.3, 1.4 and 1.2-fold IC_50_ increases, respectively, as compared to isogenic K13 wild-type parasites (**Fig. 2A**). The PB F458I K13 mutant displayed equal sensitivity to DHA as the wild-type and the UBP-1 V2721F mutant (**Fig. 2A**), in agreement with our previous observations (40). These data suggest that, despite being limited to a single cycle 24-hour exposure, the PB standard assay can distinguish even modestly, ART-resistant parasites from sensitive ones. We next investigated the DHA susceptibility of early ring-stage PB K13 mutant parasites by adapting the PF RSA (17). The PF RSA relies on exposure of early ring-stage parasites (0-3 hours post invasion) to 700nM DHA for 4-6 hours, followed by assessment of viability in the 2nd life cycle. This protocol allows drug-exposed parasites to re-invade fresh red blood cells. With this approach, current RSA parameters define *in vitro* ART resistance as survival of≥1% and ART sensitivity as <1% survival (17). Using a similar approach, we exposed ∼1.5-hour post-invasion K13 mutant PB ring-stage parasites to DHA at 700nM for 3 hours (to accommodate for the shorter life cycle in PB). Viability was assessed 24 hours later by flow cytometry-based quantification of schizont maturation and mCherry expression. Interestingly, we observed that a significant fraction of PB wild-type parasites survive exposure to DHA at 700nM, with percentage survival rates of ∼20.9% (**Fig. 2B**). This is in agreement with our previous observations that PB is less susceptible to ARTs relative to PF (40, 47). Both the UBP-1 mutant and F458I or Y505H K13 mutant parasites had the same survival rates as the wild-type line, while the M488I and R551T mutants exhibited significantly higher survival rates (32.3% or 39.0% respectively, P <0.001) (**Fig. 2B**). This is consistent with previous reports that in PF, the R539T and I543T mutations are associated with the highest rates of RSA survival (22). However, we noted inconsistencies between drug susceptibility data of the mutants in the two *in vitro* tests (standard 24-hour assay and adapted PB RSA). This might result from the inability to maintain PB in long-term culture and extend the analysis. We therefore developed a modified *in vivo* RSA, where we injected wild-type, UBP-1 V2721F, M488I and R551T parasites back into mice 24 hours after DMSO or DHA exposure in the RSA as described above, and then assessed viability by quantifying *in vivo* parasitemia on Day 4. Remarkably, percentage survival in the R551T mutant parasites significantly increased from ∼39.0% (24-hours readout) to ∼62.5%, while M488I mutant parasites survival increased from ∼32.3% (24-hours readout) to ∼38.0% (**Fig. 2C**). In contrast, the percentage survival of the wild-type and UBP-1 mutant lines did not significantly change in the extended assay, despite the minor growth defect in the UBP-1 mutant, demonstrating that the PB *in vitro* RSA and standard growth inhibition assays with 24-hour readouts may be less robust in quantifying resistance phenotypes, especially if mutant parasites are less fit (**Fig. 2C**).

**Fig. 2.**
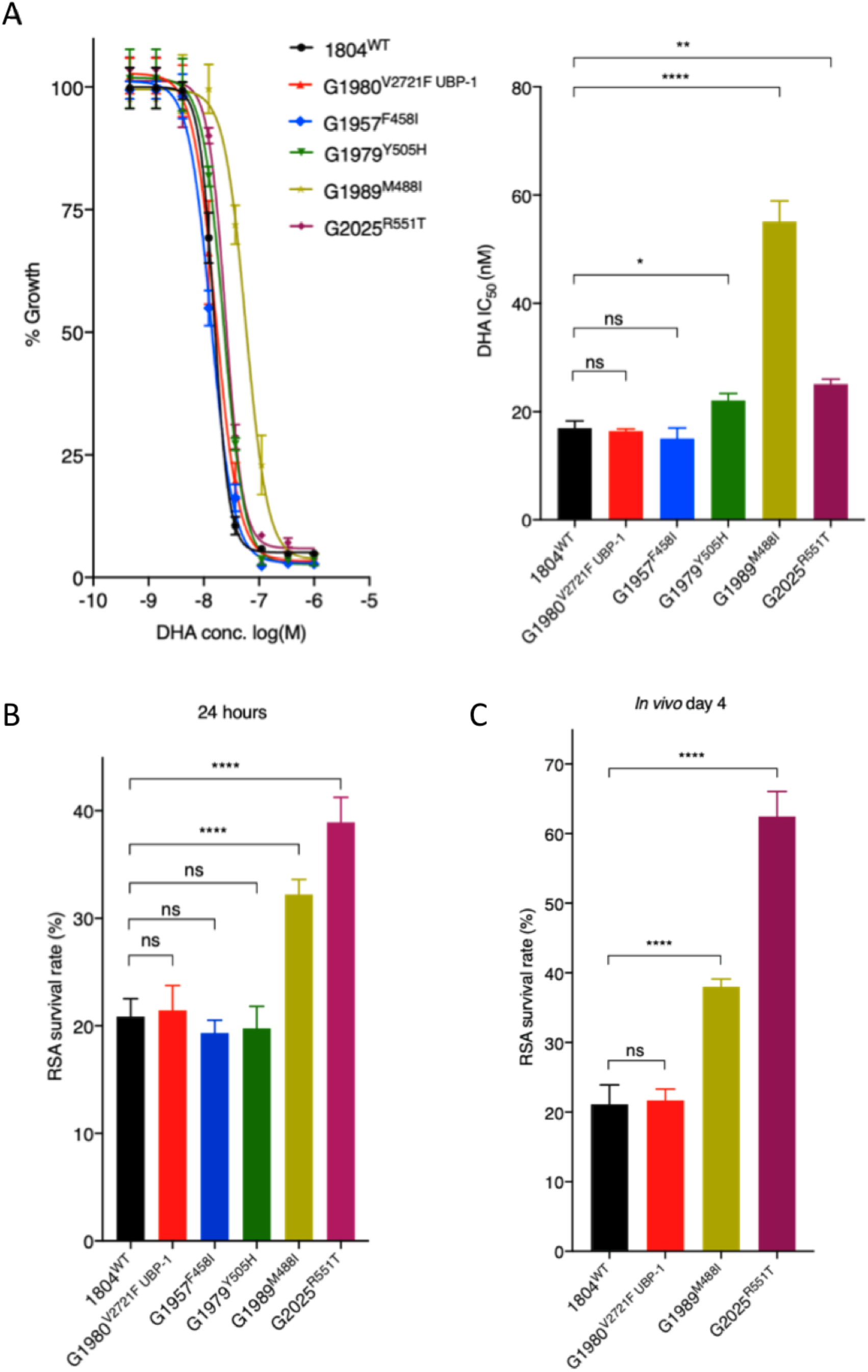
*In vitro* and *ex-vivo* susceptibility of PB K13 mutants to DHA. **A**. DHA dose-response curves and IC_50_ values for PB K13 mutant lines as compared to wild-type 1804^WT^ and the UBP-1 G1808^V2721F^ mutant line. **B**. Survival of PB K13 mutant lines in the PB RSA. Results show the percentage of synchronized early ring-stage parasites (1.5 hours post invasion) that survived a 3-hour exposure to 700nM DHA relative to DMSO-treated parasites. Survival was quantified 24 hours post treatment by flow cytometry analysis based on Hoescht 33258 DNA staining and mCherry expression. **C**. *In vivo* RSA survival for two K13 mutant lines (G1989^M488I^ and G2025^R551T^) as compared to the wild-type (1804^WT^) and UBP-1 mutant (G1980^V2721F^) controls. After *in vitro* exposure to DHA or DMSO as described above, parasites were IV injected back into mice as described in Methods. Parasitemia was quantified by flow cytometry analysis of mCherry expression on Day 4 post IV, from which % survival rates were calculated. Error bars show standard deviation calculated from three biological repeats. Statistical significance (compared to the 1804^WT^ line) was calculated using one-way ANOVA alongside the Dunnet’s multiple comparison test. Significance is indicated with asterisks; ns not significant *p < 0.05, **p < 0.01, ****p < 0.0001.

### PB K13 mutants mimic the delayed parasite clearance phenotype *in vivo* upon AS treatment and achieve faster recrudescence than wild-type parasites at high ART doses

We next investigated the *in vivo* parasite clearance rates of PB K13 mutant parasites in infected mice treated with AS. Mice were infected with a fixed inoculum of K13 and UBP-1 mutant parasites (10^5^) in four cohorts and parasitemias were allowed to rise to ∼10%. This was followed by dosing with AS at 64 mg/kg, which is slightly higher than the equivalent of the maximal human clinical dose of 4 mg/kg (mouse equivalent = 49.2 mg/kg) to accommodate for the reduced ART susceptibility observed in PB parasites. Parasitemias were quantified by flow cytometry (based on mCherry positivity) and microscopic analysis every 3 hours for the first 24 hours and at least once after the second and third doses at 24 and 48 hours respectively. Plotting parasite density in PB K13 and UBP-1 mutant parasites against time revealed that in the first 24 hours of sampling, parasite clearance kinetics did not sufficiently discriminate K13 or UBP-1 mutant parasites from wild-type. However, as the majority of dying parasites were being cleared by the host and mice received further doses, extended analysis revealed that PB M488I and R551T mutant parasites consistently and significantly persisted compared to wild-type, F458I, Y505H and UBP-1 mutant parasites (**Fig. 3A; Fig. S4**). Starting AS treatment at a high initial parasitemia (∼10%) also ensured that a good proportion of parasites would be within the early ring-stage window and therefore, would be expected to preferentially survive the first AS dose. Surviving rings could be easily distinguished as viable trophozoites at either 18, 21- or 24-hours’ time points by microscopic examination of blood smears, which enabled comparisons between parasite lines. We therefore carried out concurrent collection and analysis of thin blood smears at all time points examined for flow analysis (**Fig. 3A; Fig. S4**). Results demonstrated that enhanced survival of the first AS dose was evident for all four PB K13 mutant parasites as well as the UBP-1 mutant, compared to wild-type parasites (**Fig. S5**). Microscopy provided a more sensitive discrimination than flow cytometry-based estimation of clearance kinetics, which was unable to distinguish mutant from wild-type parasites in the first 24 hours. False positives could be due to the retention of mCherry positivity by dying parasites, as for instance, we observed that a significant proportion of wild-type parasites remained mCherry positive and were counted as viable by flow cytometry (**Fig. 3A; Fig. S4**), whereas microscopically, they were pyknotic forms (**Fig. S5A, S5G**). Remarkably, the M488I and R551T mutants remained smear positive after two consecutive AS doses (**Fig. S5E, S5F, S5K, S5L**) whereas the wild-type, F458I, Y505H and UBP-1 mutant parasites were cleared (microscopically smear negative) after 48 hours. These data suggest that the M488I and R551T mutants meet the classical definition of ART resistance as defined by the WHO, based on day 3 (second generation) microscopy positivity when accounting for the duration of the PB life cycle and the dosing intervals (4). One of the four mice in the M488I treatment group remained smear positive after three consecutive AS doses (**Fig. S5E**). These data provide evidence that PB K13 mutants modulate *in vivo* susceptibility to ARTs, resulting in a persister/delayed clearance phenotype under controlled conditions of initial parasite biomass and host immune status. Of note, we consistently used naïve mice of same age, gender, breed and genetic background.

**Fig. 3.**
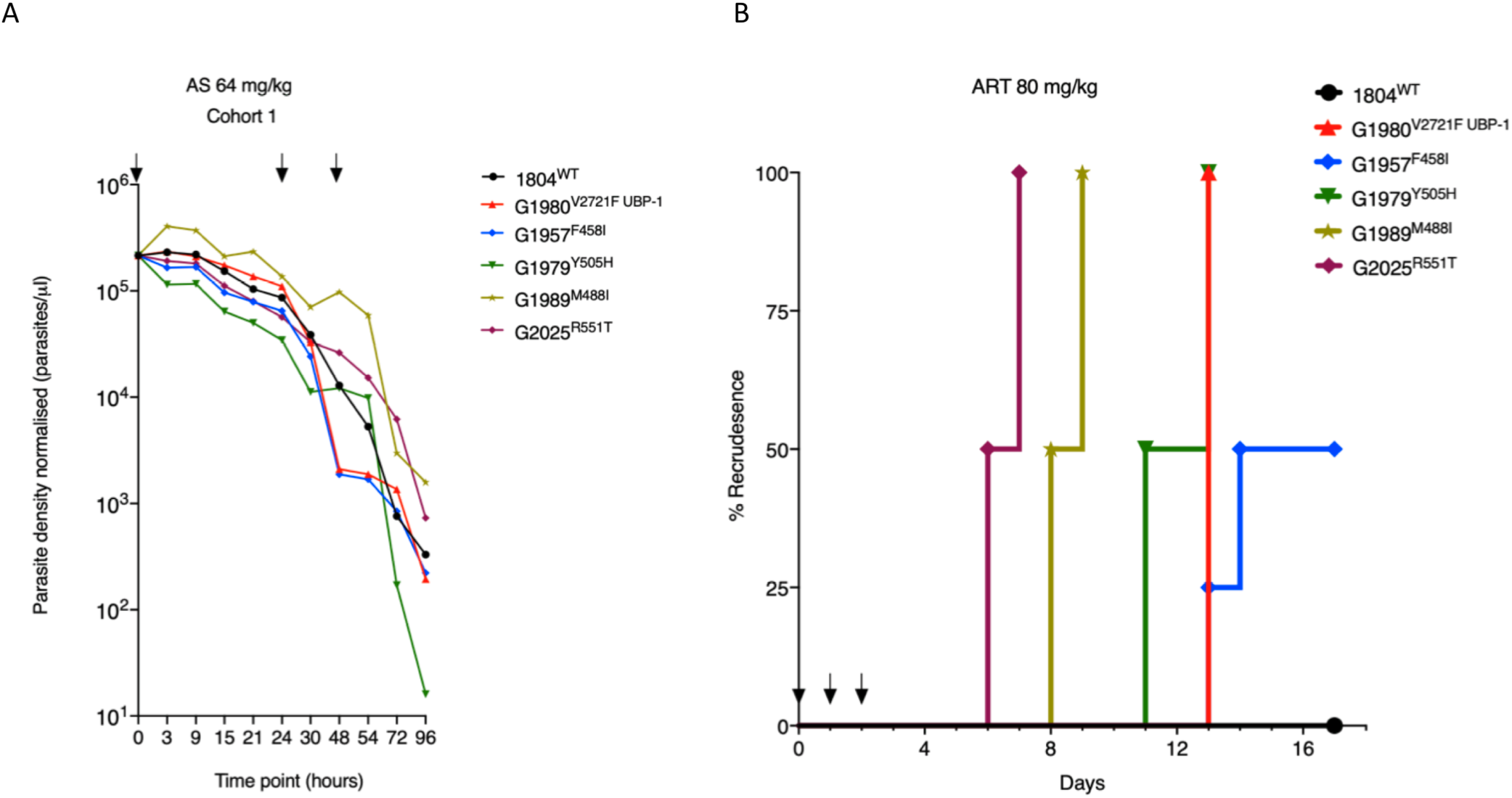
*In vivo* clearance and recrudescence rate of PB K13 mutants following treatment with AS or ART. **A**. Parasite clearance curves in mice infected with PB K13 mutant lines following treatment with AS. Six mice (in each of four cohorts) were infected with 10^5^ parasites of each of the four K13 mutants, the UBP-1 mutant and wild-type control on day 0. On day 5, at a parasitemia of between ∼10%, mice were dosed with AS at 64 mg/kg. Day 5 was the designated 0 hour timepoint for the dosing regimen. Parasite density per μl of blood was quantified based on absolute counts of mCherry-positive parasites at staggered time points for each of the two cohorts, with 5 time points in the first 24 hours (corresponding to at least 3-hour interval coverage between the two cohorts) and at least once daily thereafter. Mice were dosed three times at 0, 24 and 48 hours as indicated by arrows. Concurrent thin blood smears were prepared at each time point for microscopic analysis (Fig. S4). **B**. Kaplan-Meier plots of recrudescence in wild-type and UBP-1 mutant controls as compared to K13 mutants. A modified Peters’ 4-day suppressive test was used to monitor susceptibility of the K13 mutants to 80 mg/kg ART, a dose that effectively suppresses wild-type parasites for up to 18 days. Groups of three (UBP-1 mutant, 1804^WT^) or four mice (K13 mutants) were infected with 1x 10^6^ parasites on day 0. ART treatment was initiated ∼3 hours later and continued every 24 hours for three consecutive days (treatment days shown by arrows). Parasitemias were monitored by microscopic analysis of Giemsa-stained blood smears up to day 18 (**Table S3**). Recrudescence rates were plotted as the proportion of mice in the treatment groups that became smear positive on every individual day for the 18 days of follow-up.

Another *in vivo* marker of reduced ART susceptibility in PF is the rate of recrudescence upon AS treatment, which acts as a possible indicator of AS treatment failure. However, at pharmacologically safe doses in humans (2-4 mg/kg), ART monotherapy treatment leads to >40% recrudescence rates (1, 3), making it difficult to use this approach to separate clinically ART-sensitive from ART-resistant parasites. PB K13 mutants, therefore, provide the opportunity to test for recrudescence rates using controlled parasite inocula as well as AS or ART dose ascendency. We treated groups of mice initially infected with 10^6^ K13 mutant, ART resistant UBP-1 mutant or wild-type parasites with a daily ART dose of 80 mg/kg for three consecutive days. This ART dose sufficiently suppresses PB wild-type at equivalent parasite inocula for up to 18 days of follow-up (40). All UBP-1 mutant infections recrudesced 11 days after the last ART dose, while no recrudescence (0%) was observed for the wild-type (**Fig. 3B; Table S3**). These data are consistent with our previous observations (40). However, R551T mutant parasite infections achieved even faster recrudescence, namely 50% on day 4 after the last dosing and 100% a day later, indicating a higher level of *in vivo* resistance for this K13 mutation as compared to the UBP-1 mutant. M488I mutant parasites had a similar recrudescence profile beginning on day 6. The Y505H and F458I mutant lines both achieved recrudescence at approximately the same time as the UBP-1 mutant; however, the latter achieved only 50% recrudescence across the 18 days follow-up period (**Fig. 3B; Table S3**). These data further confirm that PB K13 mutants modulate *in vivo* susceptibility to ARTs and crucially, that recrudescence rates strongly correlate with our *in vitro* DHA RSA profiles (**Fig. 2**) as well as with *in vivo* clearance kinetics in established infections (**Fig. 3A; Figs. S4, S5**).

### PB K13 mutants are associated with an *in vivo* fitness cost but are preferentially selected for in the presence of AS or CQ

To assess the fitness of our PB K13 mutants, we performed direct head-to-head competitions with wild-type parasites under *in vivo* growth conditions. PB K13 or UBP-1 mutant lines or the parental 1804^WT^ (mCherry positive) line were mixed at a 1:1 ratios with the G159^WT^ (GFP-positive) line and injected into mice, after which changes in the proportion of GFP- or mCherry-positive parasites in the competition mixture were quantified by flow cytometry over 9 days. These assays revealed that the F458I and Y505H mutant parasites were fitness neutral relative to the G159^WT^ line, whereas the M488I and R551T mutants carried significant fitness costs (**Fig. 4A**). Both the M488I and R551T mutations were associated with high levels of reduced susceptibility to DHA *in vitro* (**Fig. 2**), delayed clearance kinetics (**Fig. 3A; Fig. S4**), and faster recrudescence following ART treatment i*n vivo* (**Fig. 3B, Table S3**). Comparatively, the R551T mutant parasites had a more severe growth defect than the M488I mutants and were completely outcompeted by the GFP-positive wild-type line by day 7 (**Fig. 4A**). This is consistent with previous observations of high *in vitro* fitness costs for the equivalent PF R539T mutation (34). In comparisons to the G159^WT^ line, the parental wild-type line (1804^WT^) was fitness neutral, whereas the UBP-1 V2721F mutant carried a minor growth defect as previously observed (40) (**Fig. S6A, S6B**). We also examined the proportion of GFP-positive versus mCherry-positive parasites over time in PB K13 mutant and wild-type parasites upon treatment with AS. Mutant parasites were mixed at 1:1 ratios with the G159^WT^ line, injected into mice and treated with AS at 50 mg/kg beginning 3 hours after infection, for three consecutive days. Monitoring of recrudescence up to day 9 revealed that, upon AS treatment, the M488I and R551T mixtures recrudesced slightly faster than the wild-type mixture and were highly enriched for the mutant population (>90%) at the time of recrudescence (**Fig. 4B**). The F458I and Y505H mutant mixtures recrudesced slightly later (**Fig. 4B**), as did the UBP-1 V2721F mutant (**Fig. S6B**) and were all significantly enriched for the mutants. In contrast, the proportions of GFP-positive versus mCherry-positive parasites in the parent 1804^WT^ and G159^WT^ competition mixture after AS treatment did not change at the time of recrudescence (**Fig. S6A**). These data show that mutant PB K13 parasites are preferentially selected for upon AS treatment, despite some carrying growth defects that rendered them at a complete competitive disadvantage in the absence of drug.

**Fig. 4.**
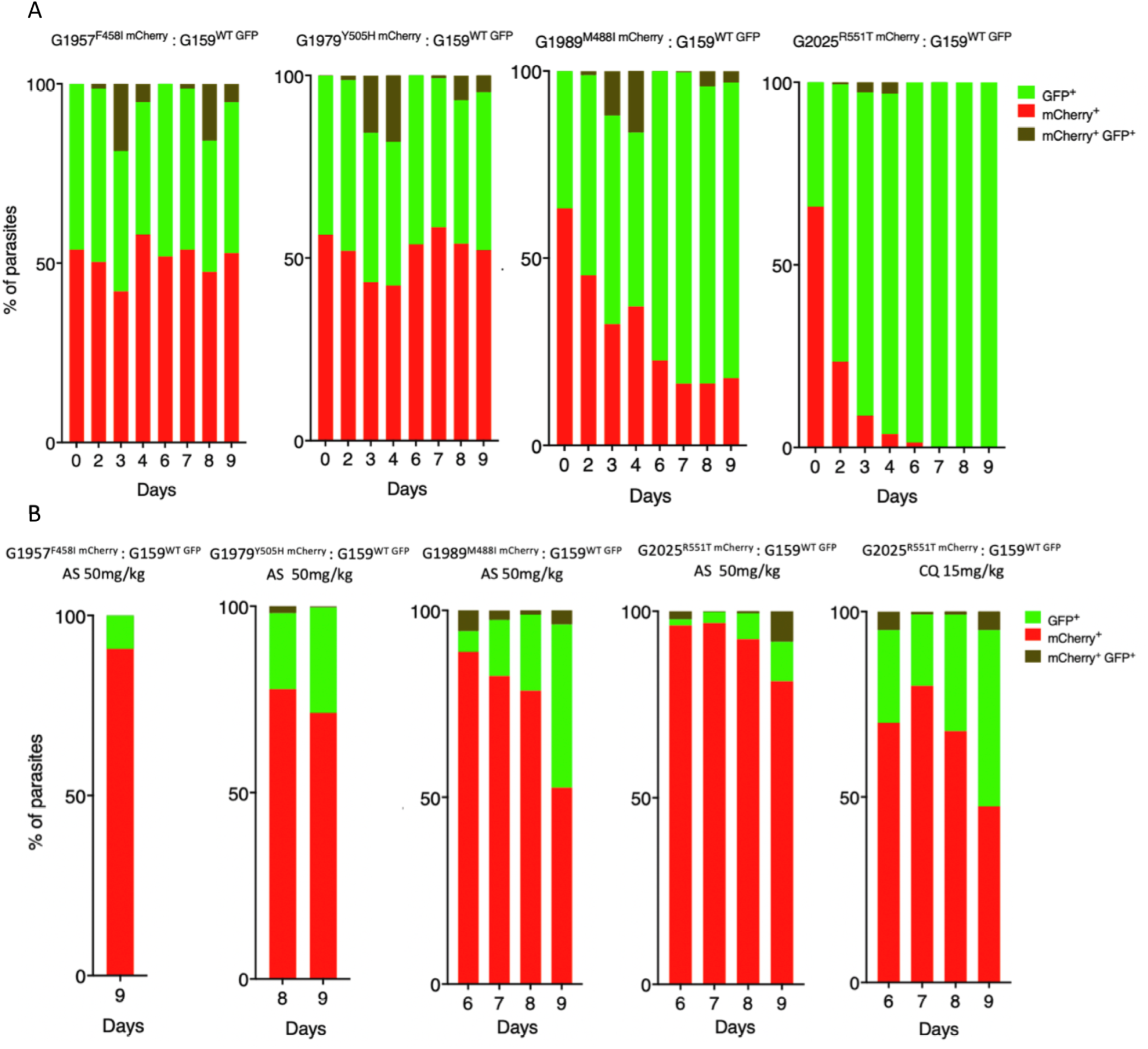
Relative fitness of PB K13 mutants in presence or absence of AS or CQ. Growth competition assays with K13 mutant lines that constitutively express mCherry as compared to the wild-type G159^WT^ line that constitutively expresses GFP in the presence or absence of drug pressure. The G159^WT^ line was mixed with a given mutant line at a 1:1 ratio in three groups of mice on Day 0. The first group was left untreated, the second group received a dose of AS at 50 mg/kg starting from 3 hours after IP injection for three consecutive doses, while the third group consisting of the 1804 ^WT^, G1980^V2721F^ and K13 mutant G2025^R551T^ lines received CQ at 15 mg/kg at similar dosing times as AS. Percentages of mCherry- or GFP-positive parasites were determined by flow cytometry as described in Methods. **A**. Percentage population changes as measured by flow cytometry of the G1957^F458I,^ G1979^Y505H^, G1989^M488I^ and G2025^R551T^ mutant lines relative to the G159^WT^ wild-type line. **B**. Proportion representation of the G159^WT^ line in mixture with G1957^F458I^, G1979^Y505H^, G1989^M488I^ and G2025^R551T^ lines on the days of recrudescence upon treatment with AS or CQ as indicated.

With the supposed role of PF K13 in mediating parasite hemoglobin endocytosis (37-39), we also speculated that PB K13 mutant parasites with strong ART resistance phenotypes might be able to modulate susceptibility to CQ (to some degree) through a similar dysregulation of the endocytic machinery. Using the *in vivo* competition assay under drug pressure as with AS above, the parental 1804^WT^ line, the UBP-1 V2721F line, and the K13 R551T mutant line were each mixed at 1:1 ratios with the G159^WT^ line and treated with CQ at 15 mg/kg. At the time of recrudescence, the proportion of 1804^WT^ parasites (mCherry-positive) did not significantly change as compared to the proportion of GFP-positive G159^WT^ parasites (**Fig. S6A**). In comparison, the UBP-1 V2721F mutant was enriched to ∼70% (**Fig. S6B**), which mirrors our previous observations that this mutation can indeed be selectively enriched by CQ (40). Interestingly, upon CQ treatment, the combination of R551T mutant parasites and the G159^WT^ line achieved recrudescence at almost the same rate as under AS pressure, with mutant parasites enriched to ∼72% (**Fig. 4B**). These data suggest that K13 mutations can also contribute to low-level protection to CQ (37, 38).

### A *Plasmodium*-selective proteasome inhibitor is potent against PB wild-type and K13 mutant parasites and synergizes DHA action

An enhanced cell stress response characterized by upregulation of genes in the unfolded protein response (UPR) is a typical signature of ART-resistant parasites (44). Resistant parasites (K13 mutants) also display enhanced activity of the ubiquitin proteasome system (UPS), a conserved eukaryotic pathway that acts downstream of the UPR by degrading unfolded proteins (43, 48). UPS inhibitors are available for cancer treatment and have been shown to synergize DHA activity in wild-type and K13 mutant PF both *in vitro* and *in vivo*, marking them as promising agents for overcoming ART resistance (43, 49). The *Plasmodium-*selective proteasome inhibitor EY5-125 is a potent antimalarial (standard IC_50_ against PF= 19nM) that acts in synergy with ART against both ART resistant and sensitive PF strains *in vitro* (50). Here, we tested the efficacy of EY5-125 against PB wild-type and K13 mutant parasites and examined its potential ability to synergize DHA action. PB wild-type and the most ART-resistant K13 mutant (R551T) parasites were found to be equally sensitive to EY5-125 (**Fig. 5A, 5B)**. Compared to PF (standard IC_50_ ∼19nM and 1hr IC_50_∼648nM), EY5-125 is much less potent in PB in both standard *in vitro* growth inhibition (IC_50_ = ∼700nM) and 3-hour assays (IC_50_ = ∼1900nM) respectively (**Fig. 5A, 5B**). These differences could be due to species-specific differences in drug sensitivity as we have observed with ARTs (40, 47) and many other drugs (51). However, combinations of DHA and EY5-125 in fixed ratio isobologram analyses revealed a strong synergistic interaction against the PB parent wild-type and K13 M488I and R551T mutant lines (**Fig. 5C, Table S4**). We also employed our *in vivo* RSA to examine whether a combination of DHA at 700nM and EY5-125 at the equivalent 3-hour IC_50_ (1.94 µM) or 2x IC_50_ (3.88 µM) could impact the parasite survival rates. Indeed, at both 3-hour IC_50_ or 2x IC_50_ concentrations, EY5-125 strongly synergized with DHA (700nM), as evidenced by significant abrogation of survival for both the wild-type and R551T mutant lines (**Fig. 5D**). These data demonstrate that proteasome inhibitors synergize DHA action in PB K13 mutants equally as well as wild-type parasites both *in vitro* and *in vivo* and have the potential to be used to overcome ART resistance.

**Fig. 5.**
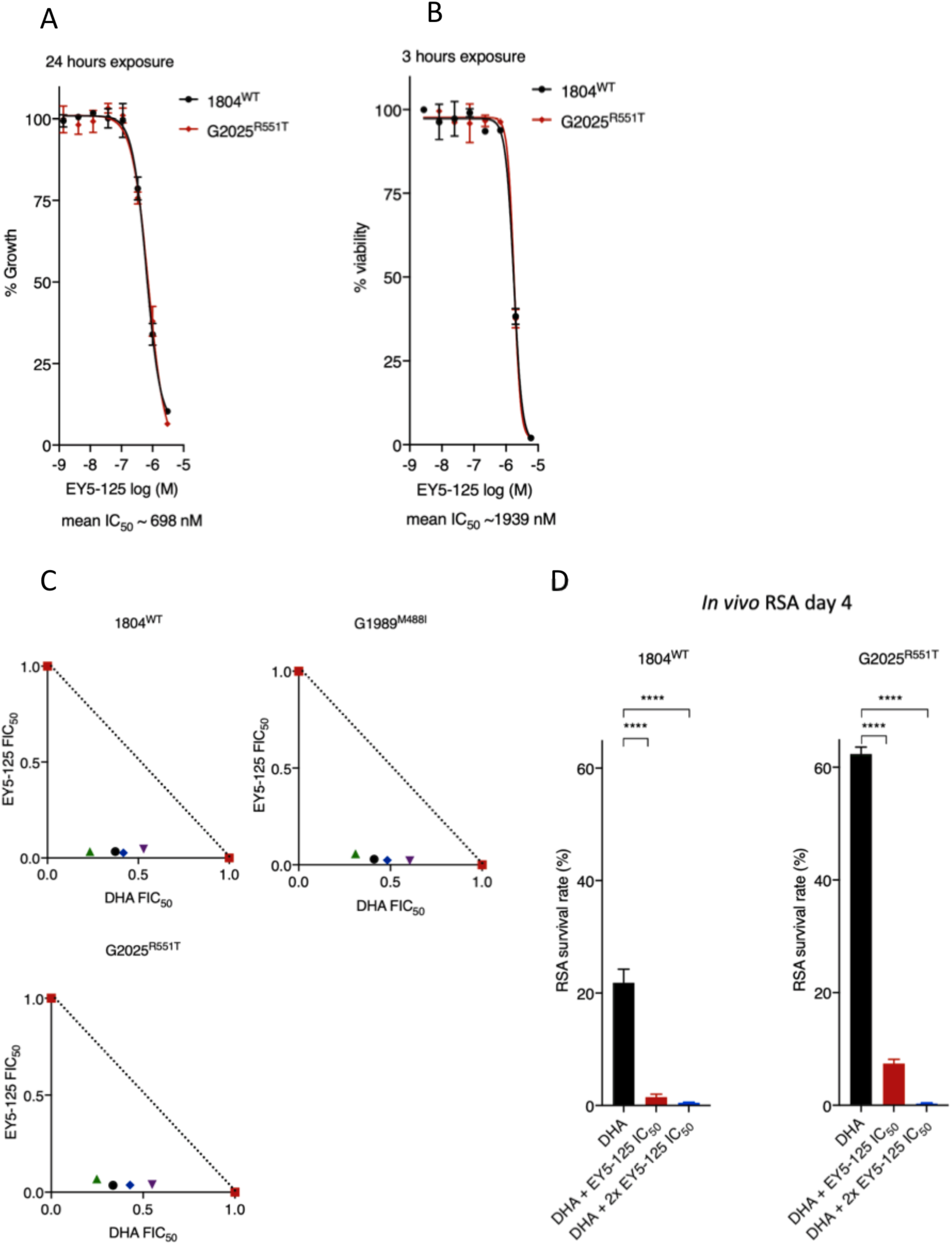
Activity and DHA synergy of proteasome inhibitor in PB K13 mutants. **A, B**. Dose-response curves and mean IC_50_ values for the *Plasmodium-*selective proteasome inhibitor EY5-125 for the wild-type 1804^WT^ and K13 mutant G2025^R551T^ lines in standard 24-hour assays (**A**) or 3-hour exposure assays conducted on early ring-stage parasites (**B**). Mean IC_50_ is a calculated average for the two lines independently screened in three biological repeats. **C**. Isobologram plots representing the interaction between DHA and EY5-125 in the wild-type 1804^WT^, G1989^M488I^ and G2025^R551T^ lines. Plots show mean FIC_50_ values (**Table S4**) for each drug calculated from three biological repeats. **D**. Synergy of EY5-125 proteasome inhibitor with DHA in the *in vivo* RSA. Parasites were exposed to DMSO or DHA at 700nM alone or in combination with EY5-125 at 3-hour IC_50_ or 2X IC_50_, then injected back into mice 24 hours later as described in Methods. Parasitemias in mice infected with drug or DMSO-treated parasites were determined by flow analysis of mCherry expression on Day 4 post IV injection and were used to calculate percent survivals relative to DMSO-treated parasites. Error bars are standard deviations from three biological repeats. Statistical significance was calculated using one-way ANOVA alongside the Dunnet’s multiple comparison test. Significance is indicated with asterisks; ****p < 0.001.

## DISCUSSION

In this study, we have successfully employed CRISPR/Cas9 editing to introduce four of the six targeted orthologous PF K13 (F446I, M476I, Y493H, R539T) mutations into the *K13* gene of the rodent model of malaria PB. Meanwhile, introduction of two mutations (C580Y and I543T) could not be achieved. As debates continue on the role of K13 in mediating *in vivo* susceptibility to ARTs (31), phenotyping of these PB K13 (F458I, M488I, Y505H and R551T) mutants provides experimental evidence for the ability of mutant K13 to confer *in vivo* resistance to ARTs in a naïve parasite genome background. These mutants displayed reduced *in vitro* susceptibility to DHA and phenocopied PF delayed clearance phenotypes upon AS treatment. Moreover, these K13 mutants achieved faster recrudescence upon ART treatment under *in vivo* growth conditions. As in PF, certain PB K13 mutations were found to cause significant growth defects, which highlights the structural and functional conservation of this protein across the two *Plasmodium* species and illustrates the fitness trade-offs that acquisition of such mutations exerts on malaria parasite physiology.

ART resistance, principally associated with mutations in K13, is now almost endemic in SEA with risks of spreading threatening the utility of ACTs that are at the forefront of malaria control programs (2). The PF C580Y K13 mutation is the most frequently observed (with >50% prevalence) and has reached fixation in most parts of SEA (19, 52). Why the PF C580Y mutation is so successful as compared to other K13 mutations remains unclear. This mutation does not associate with high RSA survival rates as compared to PF R539T or I543T mutations, nor are treatment failure rates and parasite clearance rates more significant in C580Y-harbouring parasites as compared to other K13 mutants (21, 22, 53). Do fitness constraints, founder genetic landscapes or species-specific differences between PB and PF K13 explain our failed attempts to introduce the C592Y or I555T mutations in PB? The structural homology model of the K13 propeller domain presented herein demonstrates that this region is highly conserved between PB and PF K13, with identical amino acids at the sites of mutations associated with ART resistance. Our failed attempts to introduce the PB C592Y or PB I555T mutations could therefore be more related to growth disadvantages or other deleterious effects. For example, in PF, the equivalent I543T and R539T mutations carry the most pronounced fitness costs (34) which could partly explain our inability to introduce the PB I555T mutation in PB. Moreover, PB K13 mutations were introduced into ART naïve PBANKA parasites with no history of ART exposure. These parasites might therefore be more sensitive to fitness impacts conferred by the PB I555T or PB C592Y substitution, as introduction of the equivalent PF C580Y in parasites isolated before ART was clinically introduced carried significant growth defects, as opposed to more recent Cambodian isolates where it was fitness neutral (34). A less prevalent K13 allele, PF R561H that associates with significant delays in parasite clearance and peaked in prevalence in 2012 in SEA but has since declined (53), also easily outcompeted the PF C580Y mutation in head to head competitions (36). These data suggest that acquisition and propagation of certain PF K13 alleles, mostly the C580Y substitution, require appropriate founder genome architectures to compensate for the deleterious phenotypes. In these situations, K13 mutations (PF C580Y for example), would arise in a necessary compensatory background that mitigates the deleterious growth effects leading to an initial soft sweep. In case of ACTs, these compensatory backgrounds may also serve as general templates upon which partner drug resistance mutations might arise. This seems to be the case with the recent aggressive expansion of parasite co-lineages carrying the PF C580Y mutation and piperaquine resistance determinants (9, 10).

Despite the obstacles to introducing the PB C592Y and I555T mutations, introduction of the PF R539T equivalent was achieved in PB (R551T) despite low editing efficiency in the initial transfection. We were, however, able to enrich for this mutation with AS selection applied *in vivo*, yielding almost clonal levels of the PB R551T mutant. Similar to the PF R539T mutant, clonal PB R551T mutant parasites carried the strongest DHA resistance phenotypes *in vitro* as well as the clearest AS or ART resistance profiles *in vivo*. The PF R539T and PF I543T mutations occur at relatively low frequencies in SEA with the prevalence of both mutations ranging between 0.3-3.5% (30, 35, 52). This could be due to the pronounced fitness cost of these mutations (34) limiting their expansion, which we also observed with the PB R551T mutant parasites. The combination of a naïve genomic background and species-specific differences can also be invoked to explain some phenotypic differences (growth rate and level of ART resistance) seen between mutant lines of PF and PB K13 as observed in this study. For example, PF Y493H mutants clearly associate with increased RSA survival (22, 54) and delayed parasite clearance phenotypes (35, 54, 55), unlike the PB counterpart (Y505H) that display low-level resistance to ARTs *in vitro* (in the standard assay but not in the adapted RSA) and *in vivo*. This could be due to additional underlying genetic factors in PF isolates providing an additive effect to the observed phenotypes, which would be absent in PB. Nevertheless, the other PB K13 mutations tested herein appear to directly reflect the impact of the equivalent mutations in PF. Both PB F458I (this study) and PF F446I K13 mutants are fitness neutral (56) and do not enhance RSA survival *in vitro* (56, 57), yet carry ART protective phenotypes *in vivo* (58-60). Furthermore, PB M488I K13 mutants display a significant growth defect that has not yet been characterized in the PF equivalent (M476I) and might explain its relative scarcity in SEA (61, 62).

Enhanced proteostasis is a characteristic signature of PF K13 ART-resistant parasites, which is typified by upregulation of genes in the UPR as well as enhanced activity of the UPS (43, 44, 48). Inhibition of the UPS by 26S proteasome inhibitors synergizes DHA action both *in vitro* and *in vivo*, which has offered a potential avenue to overcome ART resistance (43). Despite UPS inhibitors (which are clinically available for treatment of certain cancers) displaying activity in malaria parasites and synergizing DHA action, their translation into animal studies has been limited by host toxicity (63, 64). Recent structure-based design of *Plasmodium-*selective proteasome inhibitors has provided classes of compounds with a wider therapeutic window and improved host toxicity profiles (49, 50). These inhibitors not only display activity in diverse PF backgrounds, including those harbouring K13 mutations, but also synergize DHA action (65). Even though PB proteasome structures have not been solved, functional and life cycle conservation between this parasite and PF is pronounced. Using EY5-125, an inhibitor selective for the PF proteasome (50), we demonstrate similar activity and synergy with DHA in PB wild-type and K13 ART resistant mutants. Importantly we demonstrate these properties *in vivo*, which significantly strengthens the potential of these compounds in overcoming ART resistance in infected hosts.

In conclusion, our work provides robust experimental evidence that K13 mutations modulate *in vitro* and in *vivo* susceptibility to ARTs in the PB rodent model of malaria. The cause and effect link between PF K13 mutations with reduced ART susceptibility is strong (22, 54). However, the reason for ART clinical failure has remained obscure because, in some cases, delayed parasite clearance phenotypes have been reported in parasites carrying wild-type K13 alleles (29, 66). This lack of clarity is further compounded by a reduced correlation between K13 mutations and parasite clearance half-lives or the frequencies of recrudescence in certain cases of ART monotherapies (29). As we demonstrate in this study, some of these observations may be attributable to fitness defects in mutant parasites that could confound the interpretation of recrudescence rates. These fitness differences might be especially relevant at the relatively low ART doses used in humans, which are already known to permit higher rates of recrudescence (3). Although a recent genetic cross between a PF K13 C580Y mutant parasite and an *Aotus*-infecting K13 wild-type parasite demonstrated a lack of association of this mutation with *in vivo* ART resistance metrics (recrudescence and clearance half-life) (31), we propose that this could be due to: 1) the AS doses used being insufficiently high to clearly separate the lineages; 2) the small sample sizes used; and 3) the inherent limitation of using heterogeneous *Aotus* monkeys with varying individual histories of parasite exposure and spleen status (spleen intact or splenectomized). Our *in vitro* and *in vivo* phenotypes for the PF F446I, M476I, Y493H and R539T K13 mutation equivalents in PB support their direct involvement in mediating resistance to ARTs. Our data also provide a robust immune-replete rodent host model to test for synergistic antimalarial combinations that can restore ART efficacy and overcome resistance.

## MATERIALS AND METHODS

### CRISPR/Cas9 generation of PB K13 mutant lines

The Cas9 plasmid ABR099 was used to target mutations of interest into the PB *K13* locus (PlasmoDB gene ID: PBANKA_1356700) (40). To obtain PB equivalents of PF ART-resistant K13 mutations (PlasmoDB gene ID: PF3D7_1343700), the amino acid sequences of the two proteins were retrieved and aligned using Clustal Omega (67). To structurally align the equivalent mutations in PB K13, three-dimensional homology models of PB and PF K13 were constructed using SWISS-MODEL (PDB template 4zgc.1.A) for amino acid residues 362-738 for PB and 350-726 for PF. Models were visualized using pyMol 2.3. sgRNAs designed to target a region within 0-30 bp of the mutation of interest within the PB *K13* locus were initially cloned into the ABR099 plasmid (**Fig. S2A**). Donor repair DNA templates were designed to carry the mutation of interest in addition to silent mutations that introduced restriction sites for RFLP and that inactivated the PAMs. These templates were generated by overlap extension PCR (68) and were subsequently cloned into ABR099 plasmids carrying corresponding sgRNAs at the linker sites (**Fig. S2A**). Generated plasmids and all corresponding sgRNAs are listed **in Table S1**.

### Parasite lines and animal infections

This study employed two PB ANKA-derived parasite lines, 1804cl1 and G159. The 1804cl1 (69) and G159 (Katie Hughes, unpublished) lines express mCherry and GFP respectively, under the control of the strong constitutive *hsp70* promoter. Infections were carried out in female Theiler’s Original mice (Envigo), 6-8 weeks old, weighing 25-30 g. Infections were established either by intraperitoneal injection (IP) of ∼200 µl of cryopreserved parasite stocks, or by intravenous injections (IVs) of purified schizonts or mixed-stage parasites diluted in phosphate-buffered saline (PBS). Parasitemias in infected mice were monitored by microscopic examination of methanol-fixed thin blood smears stained with Giemsa (Sigma) or flow cytometry-based analysis of infected blood stained with Hoescht 33342 (Invitrogen). Blood from infected mice was collected by cardiac puncture under terminal anaesthesia. All animal work was performed in compliance with UK home office licensing (Project reference: P6CA91811) and ethical approval from the University of Glasgow Animal Welfare and Ethical Review Body.

### Transfections

Primary transfections were carried out in the 1804cl1 line. ∼10 µg of episomal plasmid DNA from the vectors described above (**Table S1**) was transfected by electroporation of Nycodenz-purified schizonts using the Amaxa Nucleofector Device II program U-o33, as previously described (70). Parasites were then immediately IV injected into mice. Positive selection of transfected parasites was commenced 24 hours later by adding pyrimethamine (0.07 mg/ml, Sigma) to their drinking water.

### Genotyping of transformed parasites

Parasite pellets were prepared from infected mouse blood that was lysed by resuspension in 1x E-lysis buffer (Thermo). Genomic DNA was extracted from the pellets using the Qiagen DNeasy Blood and Tissue kit according to the manufacturer’s instructions. Initial analysis of the transfected or cloned parasite lines was performed using a dual PCR-RFLP approach. PCR using primers exterior to the donor templates (**Table S1, S2**) was used to amplify fragments from the genomic DNA of the mutant lines, followed by restriction digests with the artificially introduced RFLP restriction enzymes. Relative transformation efficiencies were estimated by densitometric quantification of wild-type and mutant RFLP fragments by ImageJ2 (71). Mutations and initial RFLP analyses were further confirmed by Sanger DNA sequencing.

### Antimalarial agents

DHA (Selleckchem) at 10 mM was diluted to a working concentration in schizont culture media. The *Plasmodium-*selective proteasome inhibitor EY5-125, also known as compound 28 (50), was used to test for proteasome inhibitor synergy with DHA in K13 mutant and wild-type parasites. For *in vivo* drug treatment, AS (Sigma) was dissolved in 5% sodium bicarbonate prepared in 0.9% sodium chloride. CQ diphosphate (Sigma) was dissolved in 1x PBS. ART (Sigma) was prepared at 50 mg/ml in a 1:1 mixture of DMSO and Tween® 80 (Sigma) and diluted 10-fold in sterile distilled water immediately before administration. All drugs were prepared fresh before *in vivo* administration and drug delivery was carried out by IP injection.

### 24-hour PB *in vitro* culture and drug susceptibility assays

*In vitro* culture and drug susceptibility assays were carried out beginning with synchronized ring-stage parasites over 24-hour schizont maturation cycles, as PB can only be maintained for one intra-erythrocytic developmental cycle *in vitro*. Parasites were cultured and exposed to drugs as previously described (40), after which schizont maturation was analyzed by flow cytometry. Infected cells were stained with the DNA dye Hoechst-33258. Schizont maturation was used as a surrogate marker of growth inhibition and was quantified based on Hoechst-33258 fluorescence intensity or mCherry expression. To determine growth inhibition and calculate half-maximal inhibitory concentrations (IC_50_s), the percentage of schizonts in no-drug controls was set to 100% growth, and subsequent growth percentages in presence of drugs were calculated accordingly. Dose-response curves were plotted in GraphPad Prism.

### Adapted PB ring-stage survival assays

The PF RSA was adapted for PB to further assess the *in vitro* phenotypes of K13-mutant parasites based on a previously published protocol (17). Schizonts were obtained from *in vitro* cultured parasites as previously described (70) and injected IV into naïve mice to obtain synchronous *in vivo* infections containing >90% rings at parasitemias of 0.5-1.5%. ∼1.5 hours post injection, blood was collected from the infected mice, adjusted to 0.5% hematocrit and exposed to 700 nM DHA or 0.1% DMSO (ThermoFisher Scientific) in 96-well plates or 10 ml culture flasks. The plates and flasks were incubated with drug under standard culture conditions for 3 hours, after which the drug was washed off at least three times. Parasites were then returned to standard culture conditions in new plates and flasks with fresh schizont media for *in vitro* maturation. After 24 hours of incubation, parasite survival was assessed by flow cytometry analysis of Hoechst-33258-stained infected cells and by mCherry expression. DHA-treated samples were compared to DMSO-treated controls processed in parallel. Percent survival was calculated using the formula below:

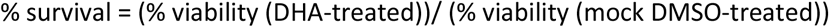

To improve the robustness of the viability readouts beyond the 24-hour flow cytometry counts, an *in vivo* expansion of the 3-hour DHA- or DMSO-exposed parasites was used for selected mutants and the wild-type control. After 24 hours of recovery, 2 ml of DHA- or DMSO-treated parasites were pelleted and resuspended in a 1ml volume, from which 200 μl was injected IV into mice. *In vivo* parasitemias were quantified on day 4 post injection, from which % survivals based on *in vivo* parasitemia (absolute counts of mCherry positive parasites) were calculated using the slightly modified formula below:

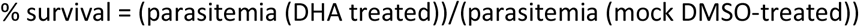

### *In vitro* isobologram drug combinations

DHA and EY5-125 drug interaction analyses in fixed ratios were carried out using a modified fixed ratio interaction assay as previously described (72). DHA and EY5-125 combinations were prepared in molar concentration combination ratios of 5:0, 4:1, 3:2, 2:3, 1:4, and 0:5 and were dispensed into 96-well plates. This was followed by a 3-fold serial dilution with pre-calculated estimates to ensure that the test wells containing the 3-hour IC_50_ of the two drugs were located near the middle of the plate. The drug combinations were then incubated with synchronized ∼1.5-hour old ring stage wild-type or K13 mutant parasites for 3 hours, after which the drugs were washed off at least 3 times. Percent viability was quantified 24 hours later by flow cytometry analysis of Hoechst 33258-stained infected cells and mCherry expression. Dose-response curves were calculated for each drug alone or in combination, from which fractional inhibitory concentrations (FIC_50_) were obtained and summed to obtain the ∑FIC_50_ using the formula below:

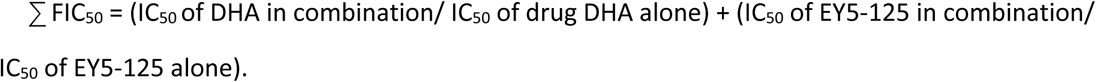

An ∑FIC_50_ of >1 was used to denote antagonism, ∑FIC_50_ <1 synergism, and ∑FIC_50_ = 1 additivity. FIC_50_ values for the drug combinations were plotted to obtain isobolograms for the drug combination ratios.

### *In vivo* drug assays: parasite clearance

Parasite clearance upon treatment with AS was used to evaluate potential delayed clearance phenotypes in K13-mutant parasites. These studies were based on a modified Rane’s curative test in established mice infections as previously described (73). Donor mice were infected with mutant lines and the wild-type control. Once a parasitemia of ∼2% was reached, blood was obtained from the donor mice and diluted in 1x PBS. 10^5^ parasites were inoculated in 4 cohorts of mice (4 mice per line) by IP on day 0 and parasitemias were allowed to rise to ∼10%, typically on day 5. On day 5, at time zero, 2μl of blood was collected and diluted 200-fold in 1x PBS. Thin blood smears were also collected at this time. All four cohorts were then dosed with AS at 64 mg/kg at 0, 24 and 48 hours. Blood sampling was performed for flow cytometry analysis and thin blood smears were prepared five times during the first 24 hours for each cohort and at least daily after that in a staggered manner that allowed for a 3-hour lifecycle coverage in the first 24 hours for at least two cohorts. Parasite density at each time point was determined by absolute cell counts and mCherry expression in 0.1 μl of whole blood diluted in PBS analyzed on a MACSQuant® Analyzer 10. Thin blood smears of parasite morphologies were analyzed by microscopy. Significant viability counts in microscopy smears were based on microscopic confirmation of at least four viable parasites in a minimum of 10 fields. Clearance kinetics of normalized parasite densities vs. time were plotted in GraphPad prism.

### *In vivo* drug assays: recrudescence

A modified Peters’ 4-day suppressive test was used to assess *in vivo* response profiles and recrudescence rates of wild-type and mutant lines as previously described (40, 74). Infections were initiated by IP inoculation of 10^6^ parasites diluted from donor mice and were followed by three daily consecutive drug doses of ART at 80 mg/kg, with the first initiated ∼3 hours post inoculation. Parasitemia was monitored by microscopic analysis of methanol-fixed Giemsa-stained smears up to day 18 or until recrudescence was observed.

### *In vivo* growth competition assays in presence or absence of drug treatment

Mutant lines in the 1804cl1 mCherry background line were mixed with the G159 GFP line at 1:1 ratios and injected by IP (total parasite inocula of 10^6^) in 3 groups of mice. The groups were left either untreated or were treated with AS at 50 mg/kg for 3 consecutive days starting 3 hours post infection, or CQ at 15 mg/kg. Parasitemias and fractions of mutant versus wild-type parasites were determined by flow cytometry-based quantification of mCherry- or GFP-positive parasite populations.

## ACKNOWLEDGMENTS

We would like to thank Diane Vaughan and the University of Glasgow flow cytometry facility for assistance. We also thank Dr. Euna Yoo (Stanford University and NCI-Frederick) for providing the EY-125 proteasome inhibitor. This work was supported in part by grants from the Wellcome Trust to A.P.W (083811/Z/07/Z; 107046/Z/15/Z and 104111/Z/14/Z). Partial funding for this work was provided by the NIH (R01 AI109023 to DAF and R33 AI127581 to MB and DAF), the Department of Defense (W81XWH-19-1-0086 to DAF) and the Columbia University - University of Glasgow Research Exchange Program. N.V.S is a Commonwealth Doctoral Scholar (MWCS-2017-789), funded by the UK government. BHS gratefully acknowledges earlier support from the Columbia University Graduate Training Program in Microbiology and Immunology (T32 AI106711, Program Director D. Fidock).

